# Simultaneous silencing of gut nucleases and a vital target gene by adult dsRNA feeding enhances RNAi efficiency and mortality in *Ceratitis capitata* adults

**DOI:** 10.1101/2024.08.01.605863

**Authors:** Gennaro Volpe, Sarah Maria Mazzucchiello, Noemi Rosati, Francesca Lucibelli, Marianna Varone, Dora Baccaro, Ilaria Mattei, Ilaria Di Lelio, Andrea Becchimanzi, Ennio Giordano, Marco Salvemini, Serena Aceto, Francesco Pennacchio, Giuseppe Saccone

## Abstract

*Ceratitis capitata*, known as Mediterranean fruit fly (Medfly), is a major dipteran pest significantly impacting fruit and vegetable farming. Currently, its control heavily relies mainly on chemical insecticides, which pose health risks and have effects on pollinators. A friendly and species-specific alternative strategy involves providing double-stranded RNA (dsRNA) through feeding to disrupt essential functions in pest insects, which is poorly explored in dipteran species. Previous reports in Orthoptera and Coleoptera species suggested that dsRNA degradation by two specific nucleases in the intestinal lumen is among the major obstacle to feeding-mediated RNAi in insects. In our study, we experimented with three-day adult feeding using a combination of dsRNA molecules that target the expression of the *ATPase* vital gene and two intestinal dsRNA nucleases. These dsRNA molecules were recently tested separately in two Tephritidae species, showing limited effectiveness [1,2]. In contrast, we observed 79% mortality over seven days, which was associated with a decrease in mRNA levels of the three targeted genes. As expected, we also observed a reduction in dsRNA degradation following RNAi against nucleases. This research illustrates the potential of utilizing molecules as pesticides to achieve mortality rates in Medfly adults by targeting crucial genes and intestinal nucleases. Furthermore, it underscores the importance of exploring RNAi-based approaches for pest management

**Simple Summary:** The control of insect pest species, mainly belonging to Orthoptera, Hemiptera, and Coleoptera species, can be based on novel emerging species-specific pesticides. These consist of dsRNA molecules delivered by feeding to insect larvae or adults, which suppress vital gene functions by RNA-RNA sequence complementarity and RNA interference. However, fewer studies of dsRNA feeding have been performed in dipteran pest insects. Two studies in Orthoptera and Coleoptera species have shown that suppressing intestinal enzymes degrading external dsRNA can improve insect mortality rates. *Ceratitis capitata* (Tephritidae), the Mediterranean fruit fly (Medfly), is a major dipteran pest significantly impacting fruit and vegetable farming. Currently, its control heavily relies mainly on chemical insecticides, which pose health risks and have effects on beneficial pollinators. Previous attempts to induce mortality by adult dsRNA-feeding in this and other Tephritidae species, such as *Bactrocera tryoni* and *B. dorsalis*, showed some effectiveness, but were often limited. We improved this method by simultaneously silencing two intestinal nucleases and a vital gene. We have found a mix of three dsRNAs able to induce much higher mortality (79%) within one week, following only three days of adult feeding.

## 1. Introduction

There are 4000 species of fruit flies worldwide, and 35% are important pests, including commercial fruits with high economic value [3]. The Mediterranean fruit fly (Medfly), *Ceratitis capitata* (Diptera: Tephritidae), is a major agricultural invasive pest that inflicts widespread damage on various fruit crops globally [3]. The females lay eggs inside ripe fruits, and the subsequent hatched larvae feed them, damaging them. It causes yearly losses of billions of euros due to decreased production, increased costs of control methods, reduced marketability of affected produce, and lost markets [4]. Traditional control methods, like chemical insecticides, face several challenges, including environmental impact (such as pollinator reduction and pollution), resistance development, and diminished effectiveness [5]. Consequently, innovative and sustainable solutions are urgently required to significantly reduce chemical pesticides in agriculture [6]. Biological control methods have been explored as a more environmentally friendly alternative to traditional insecticides for managing fruit fly populations. These methods include suppressing fruit fly populations using natural enemies, such as parasites and predators [7]. However, despite efforts in biological control, fruit fly control programs still face several challenges [3]. The Sterile Insect Technique (SIT) presents a potentially effective, eco-friendly, and species-specific control method [8]. However, its implementation is costly and challenging, mainly due to the need for extensive area-wide coordination in the suppression plan. Therefore, novel and sustainable approaches are urgently needed in parallel to or in combination with the SIT and other eco-friendly approaches.

One promising approach for pest control is using RNA interference (RNAi) to target essential genes by feeding [9,10]. Insect pests can consume artificial dsRNA molecules incorporated into their food or bait, causing a reduction in the target protein production that can affect the biological processes critical for their survival, growth, or reproduction, leading to phenotypic changes or even mortality. Over 15 years ago, Baum *et al*. [11] provided the first proof that RNAi could target specific beetle pests. Since then, numerous studies have demonstrated the effectiveness of this control strategy across various pest insect species. Environmental RNA interference (eRNAi) is emerging as a powerful tool for gene silencing, with significant potential for developing species-specific and environmentally friendly pest control methods [12,13]. Among the Tephritidae fruit flies, an adult diet, including dsRNA, induced male sterility in *B. dorsalis*, suggesting systemic RNAi [14]. The administration of dsRNA targeting genes involved in spermatogenesis, such as *boule (bol), zero population growth (zpg)*, and *doublesex* male isoform *(dsxM)*, resulted in male sterility up to 85.40%. Similarly, RNAi silencing of the sex peptide receptor mating female behavior caused female fertility in the olive fly *B. oleae* after feeding adults with transformed bacteria producing specific dsRNAs [15]. RNAi can also be effective at Tephritidae larval stages when they feed inside the fruit. However, future development of transgenic plants expressing specific dsRNA would be necessary for the delivery and sustained impact [16]. The first RNAi commercial spray pesticide (Ledprona) has been authorized by EPA this year, designed as dsRNA to target after feeding the vital gene encoding the proteasome subunit beta of the Colorado potato beetle (CPB, *Leptinotarsa decemlineata* [17]; patent WO 2020/097414).

Among potential dsRNA-based pesticides, another relevant vital target is the *vacuolar (H+)-ATPase (v-ATPase)* complex, composed of two domains (transmembrane V0 and cytoplasmic V1 domains) and responsible for transporting protons (V0) across membranes using energy from the hydrolysis of ATP (V1) [18]. V0 and V1 domains are composed, respectively, of five (a-e) and eight subunits (A-H). The *v-ATPase* complex also includes two additional accessory proteins (AC45 and M8.9). Single genes encode some subunits, while most are by multiple genes. For example, in the genome of *Drosophila melanogaster*, 33 genes encode the 15 subunits of the *V-ATPase* [19]. *D. melanogaster vacuolar ATPase* loss of function mutations causes a lethal phenotype [19,20].

Due to its wide evolutionary conservation, *V-ATPase* is a promising target for pest control development. The *vacuolar ATPase* complex is also present in the gut epithelial cells and Malpighian tubules [21], and it has demonstrated susceptibility to RNAi-based gene knockdown when orally administered in various insect species. Some species of insect pests feed on the plants’ external portions, and they can be targeted by environmental dsRNA during this life stage. Feeding larvae of major pests, including Lepidoptera, Coleoptera, and Hemiptera, with *v-ATPase*-specific dsRNAs (targeting one of the various subunits, such as *V-ATPase* A, B, D, E, or H) led to significant mortality, ranging from 40% up to 100% (see for example: [11,22–24]). In the Tephritidae *Anastrepha fraterculus*, soaking larvae (30 minutes) in dsRNA solution (500 ng/µL) targeting a chaperone protein (homolog of the human VMA21 integral membrane protein) required for proper *v-ATPase* assembly led over one week to an increase of 25% mortality with the respect of 15% mortality induced by *dsGFP* control [25]. However, insect pests of the Tephritidae family grow as larvae inside the fruit crop. Only the development of transgenic plants expressing dsRNA can solve the delivery problem at this life stage. Nevertheless, adult dsRNA feeding can also be an effective control strategy, as shown in Hemiptera, Coleoptera, and Diptera, which sometimes lead to strong mortality [26–29]. For example, targeting by RNAi the *vacuolar ATPase A subunit* (*BtvATPaseA*) in the sap-sucking pest *Bemisia tabaci* (Rhynchota), also known as white fly, caused up to 97% mortality after six days of feeding [30].

However, as the ingested dsRNA molecules are exposed to various gut enzymes, their stability and ability to reach the target cells without degradation affect the efficacy of gene silencing [31–34]. Insect dsRNA intestinal RNases seem to be naturally involved in the innate immune response against invading nucleic acids such as RNA viruses, and they can heavily limit the efficacy of orally delivered dsRNAs in inducing mortality [35–37]. Hence, research is required to explore and optimize dsRNAs’ stability in pest management applications [38,39]. Various strategies can address this issue. For example, Ortolà *et al*. [1] developed a system to produce circular dsRNA molecules in *E. coli*, making them less susceptible to nuclease degradation. They tested circular dsRNAs produced from several endogenous genes of the Mediterranean fruit fly (Medfly), including *vATPase*. Injection of circular dsRNA targeting *vATPase* (500 ng) into adult flies resulted in approximately 70% mortality within seven days, with a gene silencing of about 50-60%. Conversely, oral administration (10 adults fed with a single 10 µl drop of 1 µg/µl of dsRNA solution) led to a non-significant 15% mortality compared to the control. However, egg-laying and hatching reduction was observed.

Interestingly, the efficacy of oral delivery dsRNAs is improved when combined with the silencing of intestinal dsRNA nucleases of various pest insect species [40,41]. Tayler *et al*. [2] identified two gut dsRNA nucleases in the Australian Tephritidae *B. tryoni* and targeted them by oral RNAi together with a “reporter” gene, *yellow*. They found that co-feeding adults with three dsRNAs led to a great reduction of *yellow* mRNA levels and enzymatic function, compared with feeding *yellow* dsRNA alone.

Our study explored the effects of co-feeding with dsRNAs silencing two intestinal nucleases and a possible vital gene, a *vATPaseA* subunit. In the Medfly genome, we identified a *B. tabaci v-ATPase A* orthologue and two *B. tryoni dsRNAses* orthologues [42]. We observed that adult oral dsRNA co-feeding for three days with the three dsRNAs induced a strong reduction of target RNAs and, after six days, 79% mortality. Furthermore, we have found that in the gut juice of *C. capitata* flies fed with dsRNAs silencing intestinal nucleases, the degradation of a target dsRNAs *in vitro* is less efficient, as previously shown in *B. tryoni* by Tayler *et al*. [2]. Compared with previous studies performed in many different insect species, we achieved significantly high mortality, opening the road to the development of an effective formulation of dsRNA-biopesticide for *C. capitata*.

## 2. Materials and Methods

### 2.1. Insect rearing

The *C. capitata* strain was reared in laboratory conditions at a specific temperature (25°C) and humidity (70%) with a light:dark cycle of 12:12 hours. Adult flies are fed artificial sugar and powdered yeast extract diet in a 3:1 ratio and with water. After mating adults, females laid eggs on a vertical net, which dropped into trays containing distilled water. The embryos were collected from the water and placed on an artificial diet for larvae placed in a petri dish (400 mL distilled water, 10 mL cholesterol, 8.5 mL benzoic acid, 2.5 mL hydrochloric acid, 40 g paper, 30 g sugar, 30 g yeast powder). The third instar larvae jumped from the Petri dish and pupated in sand. We transferred the pupae to Petri dishes until their emergence.

### 2.2. RNA extraction from adult flies

Total RNA was extracted from individual flies using TRIzol™ Reagent (Thermo Fisher Scientific, Waltham, USA) according to the manufacturer’s instructions and stored at −80°C. The concentration and purity of the extracted RNA were determined by measuring the absorbance ratio at 260/280 nm using Thermo Scientific Nanodrop 2000c (Thermo Fisher Scientific, Waltham, USA). Contaminating genomic DNA was removed using an RNase-free DNase I (NEB, Ipswich, MA, USA) treatment and further controlled by RT-PCR using an intron-containing *CcSOD* gene.

### 2.3. Selection of target genes, primers design, testing and sequencing

A BLASTp search of the NCBI *C. capitata* protein Database, using the protein sequences of *B. tabaci v-ATPase A* (**[30]**; GenBank: QHB15556.1), *B. tryoni RNase1* and *RNase2* ([2]; XP_039968826.1 and XP_039967124.1) led us to select for our study orthologous proteins XP_004533376.1, XP_004530585.1, XP_004530581.1, (named respectively as *ATPaseA-CcVha68-1, CcdsRNase1* and *CcdsRNase2*) showing respectively 90%, 70% and 64% amino acid identity. Subsequently, we designed primers (see Table S1) to amplify corresponding cDNA fragments (about 500 bp long) by polymerase chain reaction (PCR) derived from the three *C. capitata* orthologous transcripts, also using the Medfly genome sequence **[42]**. We used Primer3 Input software (https://primer3.ut.ee/) with the following parameters: a length of 20 bp, a Tm of 60°C, and a similar GC content (approximately 45-55%). cDNAs were produced using LunaScript® RT SuperMix Kit (NEB, Ipswich, MA, USA). For sequencing analyses and dsRNA synthesis, the PCR was performed on cDNAs with Phusion⍰ High-Fidelity DNA polymerase (NEB, Ipswich, MA, USA). The obtained amplicons were purified using the Monarch PCR & DNA Cleanup Kit (NEB, Ipswich, MA, USA), according to the manufacturer’s instructions, and sequenced by the Sanger method (Eurofins) (dsRNase1: 613 bp, dsRNase2: 557 bp, and dsATPase: 553 bp). The subsequent alignments using “MuscleWS alignment” tool showed: 98% identity (7 mismatches, 2 gaps) to *v-ATPaseA-Ccvah68-1* (LOC101448474-XM_004533323.4) compared to the predicted sequence (Figure S1); 99% identity (1 mismatch, 0 gaps) for *CcdsRNase1* (LOC101448568-XM_004530528.2) compared to the predicted sequence (Figure S2); 99% identity (2 mismatches, 1 gap) for *CcdsRNase2* (LOC101463362-XM_004530524.3) compared to the predicted sequence (Figure S3). The sequencing analyses confirmed that the primers designed for this experiment produce amplicons whose sequences, for each target, correspond to those expected, with identity percentages around 98-99%. The differences could be due to the genetic variability of the strain we use (Benakeion) compared to sequences in the NCBI database derived from the ISPRA strain **[42]**. After the *in vitro* synthesis of dsRNA, using the MEGAscript® RNAi Kit (Thermo Fisher Scientific, Waltham, USA), according to the manufacturer’s instructions, for each selected target (*CcVha68-1, CcdsRNase1* and *CcdsRNase2*), the agarose gel showed for each dsRNAs a single band of the expected size, suggesting the absence of any off-target effects, and thus indicating their suitability for gene silencing experiments on adult individuals (Figure S4).

### 2.4. Semiquantitative analysis of gene expression

1 µg of DNase-treated total RNA for each sample (whole body, dissected head, thorax, and abdomen) was reverse transcribed using the LunaScript® RT SuperMix Kit (NEB, Ipswich, MA, USA), according to the manufacturer’s instructions. For semiquantitative analyses, the PCR was performed with One Taq 2X Master mix (NEB, Ipswich, MA, USA) on cDNA, according to the manufacturer’s instructions. cDNAs were used as templates for PCR, using specific pair oligos for each target gene (see Table S1). The individual amplicons were visualized by agarose gel electrophoresis (1.5%).

### 2.5. dsRNA feeding

Pupae were individually separated into 24-well plates until emergence, and adults emerged within 12 hours and were transferred into Petri dishes with perforations for gas exchange. A preliminary test showed that a drop of 10 µL of water-sugar (10%) solution added twice daily ensured 100% adult survival after seven days. One-day and three-day feeding experiments were conducted in parallel. We used four Petri dishes for each feeding experiment, each having 4 flies (2 males and 2 females). Four feeding/co-feeding experiments in Petri dishes were set, using drops of *dsRNases, dsATPase*, the Mix of the *dsRNases/dsATPase* (co-feeding) and the control (dsRNA-*GFP*) (see Table S3). Each dsRNA feeding (10 µL of water-sugar 10% solution; dsRNA concentration 200 ng/µL) was administered twice daily (6 hours apart). The 16 flies of the first experiment (1 day) and the 16 flies of the second parallel experiment (3 days) were sacrificed respectively in the afternoon of the second and fourth day to extract individually total RNA and perform qPCR on the four genes, including the housekeeping. A technical duplicate was performed for each RNA sample for a total of 64 qPCR reactions. The silencing effect was compared between one and three days of feeding. To evaluate the lethal effect of the *ATPase* mRNA silencing by feeding and co-feeding with dsRNAses, we conducted the experiment as previously described by using three day feeding approach (in triplicate; see Figure S7). 8 flies were equally divided into two Petri dishes (4 males in one Petri dish and 4 females in the other Petri dish), and after three days of drop-feeding, were transferred to a small cage and fed with artificial adult diet and water (see Insect rearing).

### 2.6. Real-Time Quantitative PCR (qPCR)

RNA samples were diluted to a 50 ng/µL concentration for qPCR analyses. Quantitative Real-Time analysis (QuantStudio5 Real-Time PCR system, Applied Biosystems, Carlsbad, CA, USA) was conducted to measure the impact of dsRNA treatment on the expression of investigated target genes (*CcRNase1, CcRNase2*, and *CcVha68-1*) by using specific primers listed in Table S2 and considering the housekeeping gene *RpL19* as internal reference (Table S2; [43]). A total of 256 qPCR (128 qPCR x 2 technical replicates) on the four genes and 16 negative controls were performed (see raw data supp. file). The relative expression of genes was measured by one-step qPCR, using the Power SYBR™ Green RNA-to-CT™ 1-Step kit (Applied Biosystems, Carlsbad, CA, USA), according to the manufacturer’s instructions. Gene expression data were analyzed using the 2-ΔΔCT method [44,45]. For method validation, the difference between the Ct value of one of the selected target genes and the Ct value of the reference gene *Rpl19* [ΔCt = Ct (target) - Ct (reference)] was measured against serial dilutions (400 ng, 200 ng, 100 ng, 50 ng, and 25 ng) of the extracted RNA. The equation of the obtained curve (for each target/reference pair) had a slope value of less than 0.1, indicating that primer efficiencies were approximately equal.

### 2.7. Ex vivo nuclease activity degradation assay

To assess the impact of gut nucleases on dsRNA molecules, we followed a previous protocol [2]. Eight flies (four males and four females) were fed with dsRNA-drops (*dsGFP* and dsRNases) for three days, as previously described. On the fourth day, the flies were dissected in phosphate-buffered saline (PBS 1X) to collect the gut from mouth to anus, excluding the crop and Malpighian tubules. The single dissected eight guts were placed in Eppendorf tubes and resuspended in 10 µL of PBS 1X. After a mild centrifugation step at 5000 rpm for 5 minutes to allow the release of the gut juice, the supernatants were collected and refrigerated at 4°C for 16h to allow enzymes to diffuse into the solution. Then, 100 ng of *dsATPase* was added to a diluted solution (1:20) of supernatant and incubated at RT for four time points (0, 15, 30, and 60 minutes). Samples from each time point were stored at −20°C until visualization using 1.5% agarose gel electrophoresis. Band fluorescence intensities were measured using Image Lab™ software (Bio-Rad, Hercules, California, USA).

### 2.8. Statistical analysis

The normality of the data obtained through Real-Time analysis was assessed using the Shapiro-Wilk and Kolmogorov-Smirnov tests. The expression levels of the target genes in different biological groups were analyzed using the one-way ANOVA test. When significant effects were observed (*P*-value < 0.05), Dunnett’s test was used for multiple comparisons. Survival curves of the different biological groups analyzed after dsRNA treatment were compared using the Log-rank test (Mantel-Cox). The *ex vivo* degradation assay data were analyzed using the two-way ANOVA test, as affected by dsRNA treatment and time post-treatment. All statistical analyses were performed using the software GraphPad Prism 9.0 (GraphPad Software, San Diego, USA).

## 3. Results

### 3.1. Homology search of ATPase and dsRNase orthologues in the Medfly genome and selection of three targets

Our study was inspired in part by Upadhyay *et al*. [30], which identified and targeted by oral RNAi the white fly *B. tabaci* (Rhyncota) *v-ATPase A* gene (*BtATPase*) (20 ng/µL *ad libitum* feeding for six days; 1-2 millimeter size). We have found three *C. capitata* orthologous *v-ATPase* A proteins by BLASTp analysis and selected the first hit for our study. Based on the *Drosophila* Flybase nomenclature of three related *v-ATPase* A proteins, we named this first gene *CcVha68-1* [46]. Interestingly, we have found that Ortolà *et al*. [1] targeted a *C. capitata* gene, which encodes the second hit of our BLASTp analysis (*CcVha68-2*).

The *B. tryoni* dsRNase1 (XP_039968826.1; LOC120780634; BtdsRNase1) and dsRNase2 orthologous proteins (XP_039967124.1; LOC120779055; BtdsRNase2) used in the Tayler *et al*. [2] study show 49% aa sequence identity. A BLASTp search in *C. capitata* led to finding two paralogous proteins also showing 48% aa identity (XP_004530585.1, XP_004530581.1). We concluded that they are the corresponding orthologues of the *B. tryoni* dsRNase1 and dsRNase2.

We have designed and transcribed *in vitro* 0.5 Kb long dsRNAs targeting the three gene functions. The three DNA templates for *in vitro* transcription were obtained by RT-PCR, gel-purified and sequenced. The *CcVha68-1* dsRNA sequence shows an overall 78% nucleotide identity with the paralogue *CcVha68-2*, with only two short regions of 21 bp and 68 bp having 100% identity. The two *CcdsRNases* sequences show 69-72% DNA identity over 103-137 long nucleotide (nt) regions of the corresponding paralogues, with the longest stretches being 14 nt. This suggests that the three dsRNAs (*dsATPase, dsRNase1*, and *dsRNase2*) are not expected to cause significant intergenic RNAi on the respective *C. capitata* paralogues.

### 3.2. Semiquantitative Analysis

We assessed the gene expression of the three *C. capitata* orthologues in dissected heads, thoraxes, and abdomens of adult flies by a semiquantitative RT-PCR (Figure 1). In *D. melanogaster*, based on expression data available at Flybase (https://flybase.org/, the *Vha68-1* gene is expressed in many adult body tissues, including the nervous (head), digestive (abdomen), muscle (all body, including thorax), and reproductive systems (abdomen). We have found that the *CcVha68-1* gene is expressed in all three body parts (Figure 1, **a**).

**Figure 1.**
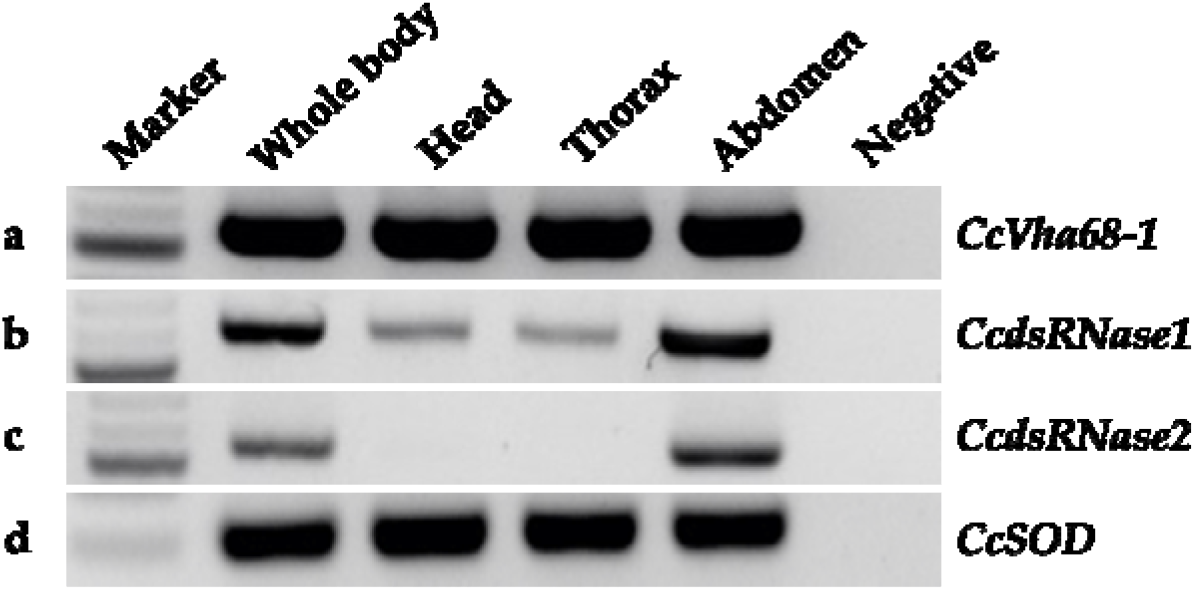
Semiquantitative analysis of *CcVha68-1, CcdsRNase1, CcdsRNase2* and *CcSOD* genes on cDNA from whole body, head, thorax, and abdomen. (**a**) *CcVha68-1* expression shows a 553 bp band in all samples; (**b**) *CcdsRNase1* expression shows a 613 bp band in all samples, with higher intensity in the abdomen; (**c**) *CcdsRNase2* expression shows a 557 bp band exclusively in the abdomen; (**d**) *CcSOD* expression (housekeeping) shows a 300 bp band in all samples (the lack of a 128bp-long intron in the amplified *CcSOD* DNA product indicted a gDNA-free cDNA sample; see Figure S5). For original agarose gels, see Figure S6.

Whilst the two *B. tryoni dsRNAses* orthologues are expressed almost exclusively in the guts of larvae and adults [2], in *C. capitata*, we found high expression of both dsRNases in the abdomen and low (*CcdsRNase1*) or no expression (*CcdsRNase2*) in the head or thorax, suggesting conservation of their gut-specific (*CcdsRNase2*) or gut-biased (*CcdsRNase1*) expression (Figure 1, **b** and **c**).

### 3.3. Gene silencing analysis

We investigated the effects of dsRNA feeding on the three targeted genes by qPCR after one day and three days of treatments. Sagri *et al*. [43] performed extensive qPCR comparative analyses of nine common housekeeping genes (HKGs) of *C. capitata* and the other Tephritidae *B. oleae* in various tissues and developmental stages. They found the *RpL19* gene to be an appropriate HKG for standardization and control reference, which we used in our study. For each of the four dsRNAs feeding experiments (*dsATPase, dsRNases*, Mix, and *dsGFP*; Table S3), we used Petri dishes, having each four flies (two males and two females) for a total of 16 flies for one day and 16 for three days of feeding. Individual RNA extractions from the 32 adult flies were performed on days two (16 flies) and four (16 flies). We conducted a technical duplicate for each of the three target genes and the reference gene and performed 128 qPCRs for one day (16 flies x four genes= 64; 64 x 2 = 128) and 128 for three days of feeding. After one and three days of feeding treatments, a reduction of 50% was observed in *CcVha68-1* mRNA levels in flies fed with dsATPase alone (Figure 2, **a** and **b**) (one-way ANOVA: *P* < 0.001). In the two one-day and three-day feeding experiments, respectively, a 60% and 70% reduction in both *CcdsRNase1* and *CcdsRNase2* mRNA levels were observed in dsRNases and Mix groups (Figure 2, **c** to **f**) (one-way ANOVA: *P* < 0.0001). Interestingly, 50% and 75% *CcVha68-1* mRNA reductions were observed after one day and after three days in flies fed with the Mix (*dsRNases+dsATPase*). Hence, after three days of co-feeding, the gene silencing of *CcVha68-1* was improved by 20-25% in the Mix group compared to dsATPase alone (Figure 2, **b**) (one-way ANOVA: *P* < 0.0001). This effect, however, is not observable when the flies are fed and analyzed after one day (Figure 2, **a**) (one-way ANOVA: *P* < 0.001). This suggests that also in *C. capitata*, as in *B. tryoni* [2], simultaneous silencing of the two intestinal nucleases (*CcdsRNase1* and *CcdsRNase2*) favors an increase in the molecular efficiency of RNAi against a third target gene, at least after three days.

**Figure 2.**
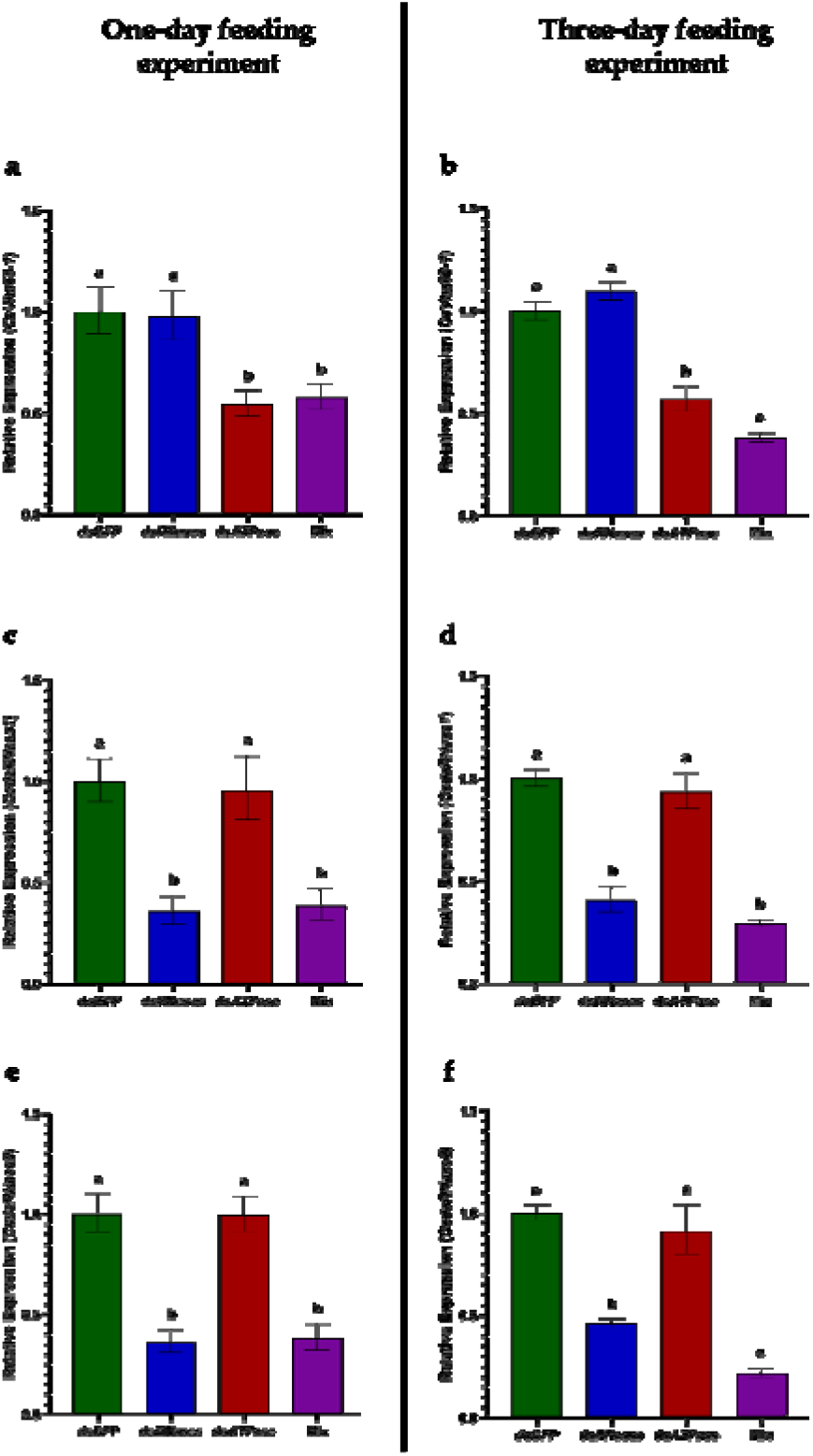
Transcripts levels of *CcVha68-1, CcdsRNase1* and *CcdsRNase2* post one-day and three-day treatments. Different letters denote a significant difference between mean values recorded for each group (the obtained values passed normality tests). The values reported are the mean ± standard error. (**a**) Relative expression of *CcVha68-1* post one-day treatment (one-way ANOVA: F(3,28) = 7.855, *P* < 0.001); (**b**) Relative expression of *CcVha68-1* post three-day treatment (one-way ANOVA: F(3,27) = 47.09, *P* < 0.0001); (**c**) Relative expression of *CcdsRNase1* post one-day treatment (one-way ANOVA: F(3,28) = 15.92, *P* < 0.0001); (**d**) Relative expression of *CcdsRNase1* post three-day treatment (one-way ANOVA: F(3,28) = 37.9, *P* < 0.0001); (**e**) Relative expression of *CcdsRNase2* post one-day treatment (one-way ANOVA: F(3,28) = 24.08, *P* < 0.0001); (**f**) Relative expression of *CcdsRNase2* post three-day treatment (one-way ANOVA: F(3,28) = 59.01, *P* < 0.0001).

### 3.4. Biopesticide activity analysis

Based on these silencing expression data, we explored the potential mortality effects after a three-day feeding experiment over seven days. Previously, Tayler *et al*. [2] targeted by dsRNA feeding the *B. tryoni yellow* gene (involved in melanization) and two *dsRNAses* and measured the melanin reduction. The simultaneous oral delivery of the three dsRNAs to groups of *B. tryoni* 10 flies for six consecutive days (one water drop of 2 µg a day) induced a very strong reduction of the *yellow* transcripts [2]. We have chosen to apply in *C. capitata* a similar strategy with some modifications. Differently from Tayler *et al*. [2], we selected a *C. capitata v-ATPase A* subunit (*CcVha68-1*) to evaluate the lethal effects of its suppression by co-feeding with dsRNAs silencing intestinal dsRNAses. We performed four parallel feeding/co-feeding dsRNA experiments, each including dsRNAs from respectively 1) *CcVha68-1*, 2) *CcdsRNAse1* and *CcdsRNAse2*, 3) the previous three genes for co-feeding (Mix), and 4) from *GFP* as a negative control (dsATPase, dsRNAses, Mix, and *dsGFP*) (Table S3). We fed/co-fed groups of eight newly emerged adult flies, with each of the four dsRNA samples, for a total of 32 flies. The eight flies were divided into two subgroups in two Petri dishes (two males and two females for each; hence, 32 flies were divided into eight Petri dishes). We performed this experiment in three biological replicates, including 96 flies in 24 Petri dishes (32 flies x 3 replicates; Figure S7, columns of the three replicate experiments). We provided to each Petri dish, twice a day (six hours apart), a 10 µL drop of water-sugar 10% solution containing 200 ng/µL dsRNA. Visual inspection showed that all four flies in each Petri dish simultaneously consumed the water drop in 10-15 minutes. By this approach, we can deduce that, assuming an equivalent feeding ability, on average, each of the four flies ingested 5 µL of solution a day (2.5 µL in the morning and 2.5 µL in the afternoon), which contained 1µg of dsRNA (on average, 3 µg for three days for each fly).

We compared mortality rates at days five and seven following the first three days of drop-feeding with the various dsRNA solutions (Figure 3; supp. raw data). *dsGFP* had no substantial lethal effect on day 5 (1/24 died; 4%) or day seven (2/24 died; 8%) (Figure 3, green line). Similarly, dsRNases feeding led to 8% (2/24) and 16% (4/24) mortality rates (Figure 3, blue line). dsATPase induced a higher mortality of 29% (7/24) and 37% (9/24), respectively (Figure 3, red line). The Mix (co-feeding with *dsRNases+dsATPAse*) was much more effective, inducing a mortality of 46% (11/24) and 79% (19/24) respectively (Figure 3, violet line). If we add the mortality percentages at day seven of the two single treatments with dsRNAses (17%) and dsATPase (37%), we obtain a 53% mortality. The observed 79% mortality with the dsRNA Mix suggests its synergistic rather than additive lethal effect. Comparing the triplicates, the Mix induced the death of 8/8, 6/8, and 5/8 flies respectively (the first two experiments were carried out with the same dsRNA batch; supp. raw data). Based on our and previous data [2], we speculated that the silencing of gut nucleases prevented, at least partially, the degradation of the dsRNA *ATPase* (*CcVha68-1*) molecules and boosted its silencing action on the endogenous vital function.

**Figure 3.**
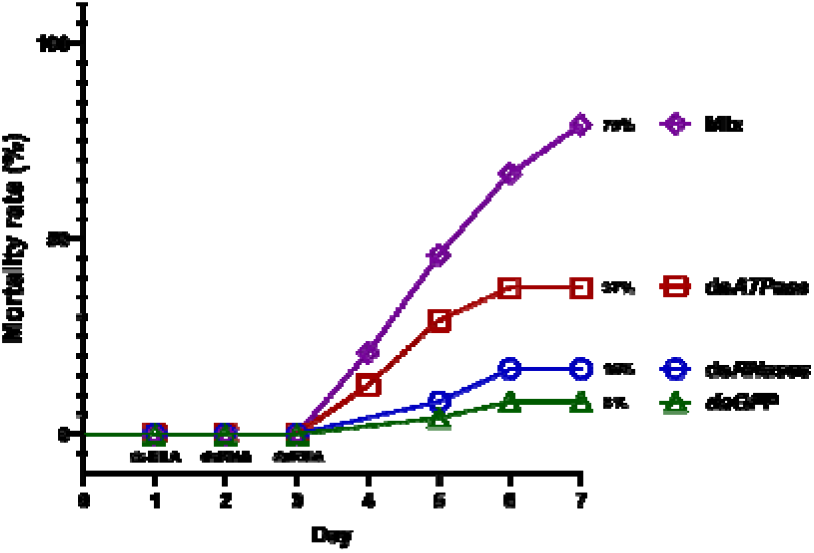
The mortality rate of treated and control flies. (Log-rank (Mantel-Cox) test: □2 = 34.53; df = 3; P < 0.0001). The feeding with dsRNAs was performed during the first three days. The mortality percentages are referred to day seven only.

To further control the molecular efficacy of *CcVha68-1* silencing in the four dsRNA feeding groups, we repeated, in parallel to the mortality experiments, the qPCR on an additional 16 flies (sacrificed at day four) in biological duplicates (a total of 32 flies; technical duplicates = 64 qPCR). As in the previous qPCR, we confirmed a *CcVha68-1* mRNA reduction of 75% (Figure S8, 4th column) (one-way ANOVA: *P* < 0.0001).

### 3.5. Nucleases activity ex vivo assay

Tayler *et al*. [2] showed in *B. tryoni* that silencing of the two intestinal *dsRNAses* by adult injection of the corresponding dsRNAs protected in an *ex vivo* assay from degradation of a target dsRNA (different from the *yellow* dsRNA used in the main experi ent) added into a test tube containing dissected guts. We preferred feeding adult flies with dsRNAs rather than injecting them and monitoring *CcVha68-1* dsRNA degradation to simulate more realistic conditions. After three days of dsRNAses or *dsGFP* feedings (control), RNA nuclease activities present in the one gut juice content of eight dissected adults (eight biological replicates for dsRNAses feeding and eight biological replicates for *dsGFP* feeding) were assessed in *ex vivo* assay, adding *in vitro CcVha68-1* dsRNAs (dsATPase; 100 ng). We monitored its degradation at four intervals (0, 15, 30, and 60 minutes; Figure 4, **a** and **b**). The dsRNA degradation was assessed by pixel intensity-based analyses after an agarose gel electrophoresis run (Figure 4, **b**). In all replicates, the dsATPase was degraded entirely (100%) within 60 minutes when incubated in the gut juice of dissected *dsGFP*-fed adults, indicating no protection from degradation. In contrast, in the dsRNases group, dsATPase was partially degraded (about 70% of deg adation) within the same time interval (two-way ANOVA: *P* < 0.0001), indicating a 30% protection efficacy (Figure 4, **b**). At 30 minutes, in the dsRNases-fed adult guts, the protection efficacy of dsATPase is 70%, while in the *dsGFP*-fed, only 20% (Figure 4, **b**). These data suggest that the gene silencing of both gut nucleases by co-feeding significantly slows down the *CcVha68-1* dsRNA degradation *in vivo*, improving both the *CcVha68-1* mRNA reduction and the related lethal effect.

**Figure 4.**
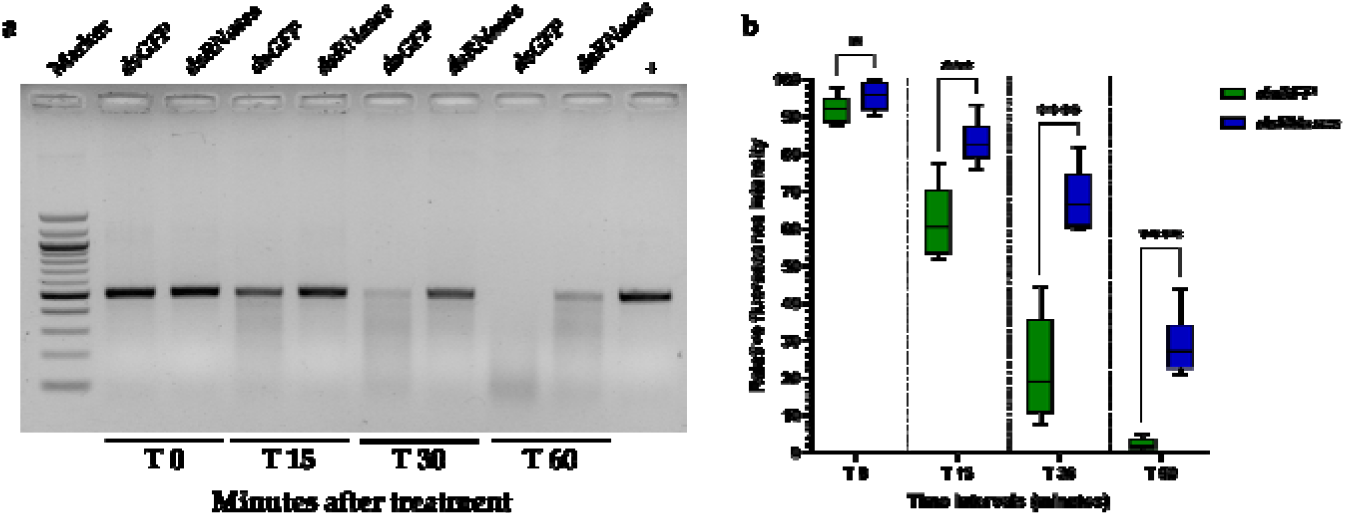
*Ex vivo* degradation assay of dsATPase molecules (100 ng for each sample). (**a**) An example of one gel electrophoresis analysis of dsRNA degradation at different time points (0, 15, 30, and 60 minutes) after gut juice (*dsGFP* and dsRNases groups) treatment (see Figure S9 for all agarose gels). The last lane represents positive control of dsRNA with no gut juice. (**b**) Overall trend (n=8 for each group) of nuclease degradation activity at different time intervals after treatment. Asterisks denote a significant difference between mean values recorded for each group (the obtained values passed normality tests). The values reported are the mean ± standard error. Both the gut juice treatment (two-way ANOVA: F(1,14) = 74.34, P < 0.0001) and the time post-treatment (two-way ANOVA: F(2.080,29.12) = 554.2, *P* < 0.0001) significantly affected the degradation of dsATPase.

## 4. Discussion

Climate change poses new risks of alien pest insect invasion and expansion into new territories [47]. Furthermore, insecticide resistance is becoming a serious problem also in *C. capitata* [48,49]. The European Green Deal has as a milestone the so-called Farm to Fork Strategy, aiming to develop novel food systems that are fair, healthy, and environmentally friendly (https://food.ec.europa.eu/horizontal-topics/farm-fork-strategy_en). This strategy sets out a key objective: a 50% reduction by 2030 in the use and risk of chemical pesticides. The development of novel solutions, including biopesticides based on dsRNAs, is urgent but limited in many cases. It is also due to the identification of very effective vital gene targets and the degradation of dsRNAs in the insect gut. The limitation of the dsRNA production costs seems to have been overcome recently, as commercial dsRNA products such as Ledprona (dsRNA specific for a proteasome subunit of Colorado potato beetle *L. decemlineata*; EPA-registered, USA) and the Ledprona-based Calantha spray-formulation (EPA-submitted, USA) [50] are close to reaching the market.

Searching for vital gene targets expressed in insect guts in the last few years led to the identification of some candidates. dsRNA feeding to target these genes led to a high degree of mortality only in a few cases, including, for example, one hemipteran (*B. tabaci*), and three coleopteran species (*Diabrotica virgifera virgifera, L. decemlineata*, and *Anthonomus grandis*). Upadhyay *et al*. [30] caused 97% mortality in *B. tabaci* (Hemiptera) adult white flies (piercing-sucking mouth parts adapted for feeding on plant sap) after six days of *ad libitum* dsRNA *ATPase A* (*Drosophila Vha68-1* orthologue; see results) feeding with liquid diet (20 ng/µL; adult size is small, only 1-2 millimeters). Rangasamy and Siegfried [26] fed the coleopteran *D. virgifera virgifera* (chewing mouthparts adapted for biting and grinding solid plant tissues) with an artificial semi-solid *ATPase* (we have found it corresponds to the *Drosophila Vha68-2* orthologue; data not shown) dsRNA diet (diet plug 4 mm; 2 mm for each adult, containing 1000 ng dsRNA every three days), causing 95% mortality after 2 weeks. On the contrary, in the Tephritidae *B. dorsalis*, 14 days long continuous feeding of dsRNA or dsRNA-expressing bacteria to target *v-ATPase D* induced only a 30% mRNA decrease but no mortality [51]. In the mosquitoes, *Aedes aegypti* oral delivery of *v-ATPase A* (we found by BLAST that it is the orthologue of *Drosophila Vha68-2*; data not shown) dsRNA (10 mosquitoes fed with 1000 ng/µL 10% sucrose *ad libitum* for 24h) failed to induce mortality after 48h. Ortolà *et al*. [1] used novel circular dsRNAs to target *v-ATPase* (*Vha68-2* orthologue). Injection of circular dsRNA (500 ng) into adult flies resulted in approximately 70% mortality within one week, with a molecular gene silencing of about 50-60%. Conversely, oral administration (20 adults in triplicates fed with a single 10 µl droplet of 1 µg/µL of dsRNA solution 30% sucrose for three days; theoretically, each fly ingested a total of 1.5 µg in three days) led to non-significant mortality (15%) compared to the control, but to a 48% reduction of mRNA levels. In summary, the mortality rates induced by feeding with dsRNA targeting *v-ATPase* subunits can vary dramatically (0-97%), generally higher in Hemiptera and Coleoptera than in Diptera.

We selected the *v-ATPase Drosophila Vha68-1* orthologue in *Ceratitis capitata* as a target for our study. When we fed for three days *C. capitata* adults with dsRNAs targeting only *CcVha68-1* (expected on average 3 µg ingested dsRNA for each fly), we observed over one week a 37% mortality rate, in contrast to the non-significant mortality observed by Ortolà *et al*. [1]. This difference could be due to different reasons: 1) the targeting of different *v-ATPase A* genes, 2) use of circular versus linear dsRNAs, 3) the amount of delivered dsRNA, 4) differences in the strains of *C. capitata* (it would be a useful strategy also to exchange *C. capitata* strains among the two laboratories), 5) the oral administration protocol: we fed four flies, twice a day for three days with 10 µl droplet (200 ng/µL dsRNA); Ortolà *et al*. [1] used 20 flies and once a day 10 µl droplet (1 µg/µL dsRNA) for three days. It would be interesting to test, for example, circular dsRNA targeting *CcVha68-1* instead of *Vha68-2* to investigate if the lack of mortality was due to a different target. For example, Taning *et al*. [13] found that feeding of *D. suzuki* adult flies with dsRNAs targeting the *Vha26* subunit induced very low mortality values.

Interestingly, Whyard *et al*. [52] targeted by dsRNA oral feeding (up to 3 µg/µL, 20% sucrose *ad libitum* for one week) the E-subunit of the *v-ATPase* gene in a broader range of insects, including larvae of a beetle (*T. castaneum*), moth larvae (*M. sexta*), aphid nymphs (*A. pisum*), and dipteran larvae (*D. melanogaster*; these larvae were soaked with liposome-dsRNA for only 2 hours). These authors observed a 50-75% mortality rate over one week. It will be interesting to investigate if *v-ATPase E*-subunit dsRNA feeding has a similar high efficacy in adults of *D. melanogaster* and of other dipteran species, including *C. capitata*, as well as to combine the dsRNA protection also with nuclease silencing.

Protection of orally supplied dsRNA can also be achieved by suppressing the degrading activities of intestinal nucleases which are among the major causes leading to differential RNAi efficiency reported among insects [36]. Tayler *et al*. [2] delivered dsRNAs (1 µg) targeting *dsRNase-1* and *-2* into adults of the other Tephritidae *B. tryoni* by injections because RNAi is more efficient than by feeding and performed an *ex vivo* assay. The three gut juices from the untreated flies and injected flies digested 100% and 30% of a target dsRNA (750 ng) *in vitro* in 1h, revealing a 70% increase in the protection from dsRNA degradation.

Within the *C. capitata* genome, we have found two nucleases orthologous to those described in *B. tryoni* and other species [2]. As in *B. tryoni*, the *CcdsRNase1* and *CcdsRNase2* seem to be expressed either predominantly (*CcdsRNase1*) or exclusively (*CcdsRNase2*) in the abdominal region (likely the guts). We performed in *C. capitata* an experiment similar to Tayler *et al*. [2] with two modifications: we fed *C. capitata* adult flies with dsRNases, instead of injecting them, and we targeted a third gene, a vital one. Our *ex vivo* assay showed that the *C. capitata* gut juices from untreated flies and dsRNases-fed flies digested 100% and 70% of 100 ng of dsRNA in 1 h (in a test tube). This data showed that the co-silencing of two nucleases by dsRNA feeding increased by 30%, the protection of dsRNA from degradation. The lower efficiency in protection with the respect of Tayler *et al*. [2] study is likely due to increased RNAi efficiency after the injection rather than feeding.

A second analysis that can be performed to investigate the effect of the nucleases silencing is to measure by qPCR the mRNA levels of a third target gene. Tayler *et al*. [2] co-fed for six days with a dsRNAs mix (2 dsRNases + dsRNA-*yellow*), a group of ten flies and induced a 100% reduction of the *yellow* RNA levels compared to *GFP* dsRNA feeding. Oral feeding with only dsRNA-*yellow* induced a 70% reduction (see Figure 3, C in [2]). Our data showed that feeding with only dsATPase induced a 50% reduction of *CcVha68-1* mRNA levels, while co-feeding of dsRNA Mix (2 *dsRNases + dsATPase*) a 75% reduction of *CcVha68-1* mRNA levels was observed (compared to *GFP* dsRNA feeding). In both studies, we can conclude that the co-silencing of intestinal nucleases improved by 25-30% the molecular silencing of a third gene (*yellow* and *CcVha68-1*).

The final experiment of our study was to measure the mortality rate induced by co-feeding with dsRNAs silencing the two dsRNases and the *ATPase A* subunit (*CcVha68-1*). We decided to perform a three days co-feeding experiment with only two droplets of 10 µL every day (200 ng/µL) to maintain lower the potential future costs of a biopesticide application, instead of an *ad libitum* feeding during a 1-2 week period used in other studies, targeting only the *v-ATPase* gene function [26,30,51,52]. Similarly, Ortolà *et al*. [1] fed *C. capitata* flies once a day for three days, monitoring the mortality over one week.

Our data showed that the co-silencing of the three genes by adult feeding, compared to silencing of only *CcVha68-1*, improved the mortality rate by 42% (from 37% to 79%) in a week period. This observation is coherent with the previous *ex vivo* experiment in which nucleases silencing protected from degradation the target dsRNA (an increase of 30% after 1h) and with the reduction of 75% of the *CcVha68-1* mRNA,

Our data offer further confirmation that intestinal nuclease activity reduces the efficacy of oral RNAi [2,32,36,38,53–56]. A simple addition of the percentages of mortality at day seven of the two single treatments with dsRNase (17%) and dsATPase (37%), leads to a 53% mortality. The 79% mortality we observed, using the dsRNA Mix suggests a synergistic rather than additive lethal effect of the co-feeding, and that dsRNases co-silencing improved the mortality rate of 25%.

Also, Spit *et al*. [40] silenced by oral feeding simultaneously in the coleopteran *L. decemlineata* (potato pest; Colorado Potato Beetle, CPB; chewing mouthparts to feed on solid plant tissues) two intestinal dsRNases and a vital gene (the authors failed to provide DNA sequence of this gene, named as *Ld_lethtgt*). They fed pre-pupal CPB with one dose of 1400 ng of dsRNases1/2 and repeated the single dose feeding post-emergence in the adults, adding 500 ng dsRNA of *Ld_lethtgt* vital gene. They observed that after eight days, there was 82% mortality. The adults fed with a single dose of dsRNA (500 ng) targeting only the vital gene led to 75% mortality. Hence, compared to our data, they observed only a mild improvement in mortality (7%) when co-silencing also the intestinal nucleases.

Similarly, Almeida Garcia *et al*. [38] micro-injected into young adults of the cotton boll weevil (*A. grandis*; Coleoptera; piercing-sucking mouth) dsRNAs targeting three dsRNA nucleases (500 ng for each in a sucrose 5% water drop) and after two days of starvation fed them with dsRNA targeting the vital *Chitin synthase II* (500 ng in a sucrose 5% water drop) (unfortunately, these authors missed performing a final co-feeding experiment avoiding microinjections, to render their method more applicable in the field). As they observed after 10 days that the nuclease suppression (injection at adult stages) led to 15% mortality, while the *Chitin synthase II* dsRNA feeding led to 60% mortality, the combined mortality of single dsRNA treatments was 75%. The application of both silencing led to an increase of only 10%, reaching 85% in mortality, while we observed a 25% increase in our experiments.

The cause of death in insects following oral delivery of dsRNA targeting *ATPase* could be due to the critical reduction in ATP production, leading to cellular energy failure. The latency in the onset of mortality (few days) can be explained, for example, by the time required for the RNAi process to effectively reduce *ATPase* levels, by the gradual depletion of ATP, by the individual variability in gene knockdown efficiency/speed, and by possible compensatory mechanisms within the insect’s cells.

## 5. Conclusions

We conclude that our feeding RNAi strategy achieved a higher and faster mortality rate for the first time in a Tephritidae species, close to those achieved only in coleopteran and hemipteran species [11,30]. We will explore how to achieve a 100% mortality rate shortly. We will modify some of the parameters of the applied method: 1) extending the feeding time, 2) increasing dsRNA quantity and concentration, 3) use of nanoparticles and liposomes, either as substitute of dsRNases silencing or in conjunction with them, 4) selection of novel intestinal vital target genes.

It is desirable to develop a standard protocol of oral RNAi to be applied as a control reference, to better compare different studies in the insect RNAi community. For example, in our protocol, we can measure approximately the quantity ingested by each fly and the time required for ingestion. In parallel, a second protocol could be designed which mimics more real field applications, once good dsRNA combinations and delivery will be found. Furthermore, improved nomenclature and gene models of the promising *ATPase A* subunits and *dsRNases* in the various Tephritidae species, including paralogues and phylogenetic comparisons, are required to better define the chosen paralogue targets in future studies (Volpe *et al*., manuscripts in prep.).

## Supplementary Materials

The following supporting information can be downloaded at: www.mdpi.com/xxx/s1, Figure S1: Alignment between dsRNA sequence of *v-ATPase A* (*CcVha68-1*) gene and XM_004533323.4 (NCBI); Figure S2: Alignment between dsRNA sequence of *RNase1* (*CcdsRNase1*) gene and XM_004530528.2 (NCBI); Figure S3: Alignment between dsRNA sequence of *RNase2* (*CcdsRNase2*) gene and XM_004530524.3 (NCBI); Figure S4: Agarose gel of the synthesized dsRNAs: dsRNase1(613 bp), dsRNase2 (557 bp), and dsATPase (553 bp); Figure S5: Agarose gel of *CcSOD* gene (housekeeping) expression in different adult tissues; Figure S6: Agarose gel of *CcVha68-1, CcdsRNase1*, and *CcdsRNase2* gene expression in different adult tissues; Figure S7: Schematic illustration of the mortality assay experiment; Figure S8: Transcript levels of *CcVha68-1* gene after three-day feeding experiment; Figure S9: Agarose gels of *ex vivo* experiment; Table S1: Primers for *CcSOD* gene amplification and the synthesis of dsRNAs target; Table S2: Primers for qPCR analysis; Table S3: dsRNAs feeding experimental groups.

## Author Contributions

Conceptualization, G.S. and G.V.; methodology, G.V., I.D., A.B., F.P.; validation, S.A., M. S. and F.P.; formal analysis, G.V..; investigation, G.V., S.M.M., N. R., F. L., M. V., D. B., I. M.,; resources, G.S..; data curation, G.V., S. A.; writing—original draft preparation, G.S., G.V., writing—review and editing, G.S., G.V., E.G.; visualization, G.S, G.V., S.A.; supervision, G.S.; project administration, G.V.; funding acquisition, G.S. All authors have read and agreed to the published version of the manuscript.

## Data Availability Statement

The authors confirm that the data supporting the findings of this study are available within the article and its supplementary materials.

## Acknowledgments

We thank the following Master thesis students Fulvio Bertolotto, Domenico De Falco, Stefania Liguori, Michela Mazzeo, David Torrente for their technical support in rearing the insects used in this study. We thank the Model Organisms Core and the Computational Biology Core of the Department of Biology for their support in this research.

## Disclaimer/Publisher’s Note

The statements, opinions and data contained in all publications are solely those of the individual author(s) and contributor(s) and not of MDPI and/or the editor(s). MDPI and/or the editor(s) disclaim responsibility for any injury to people or property resulting from any ideas, methods, instructions or products referred to in the content.

