## Supplementary materials for "Simultaneous silencing of gut nucleases and a vital target gene by adult dsRNA feeding enhances RNAi efficiency and mortality in *Ceratitis capitata* adults"

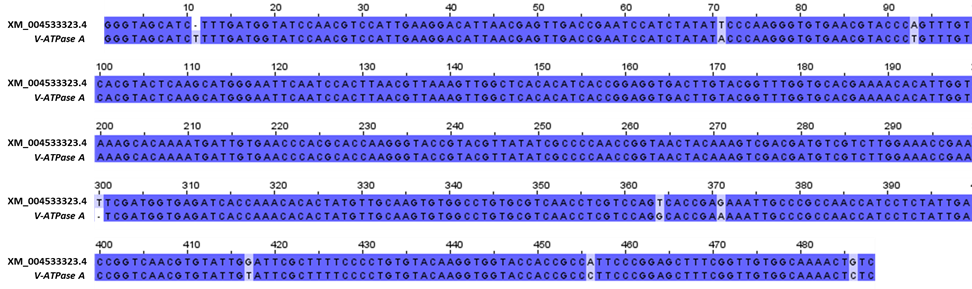

**Figure S1.** Alignment between dsRNA sequence of *v-ATPase A (CcVha68-1)* gene and XM_004533323.4 (NCBI).

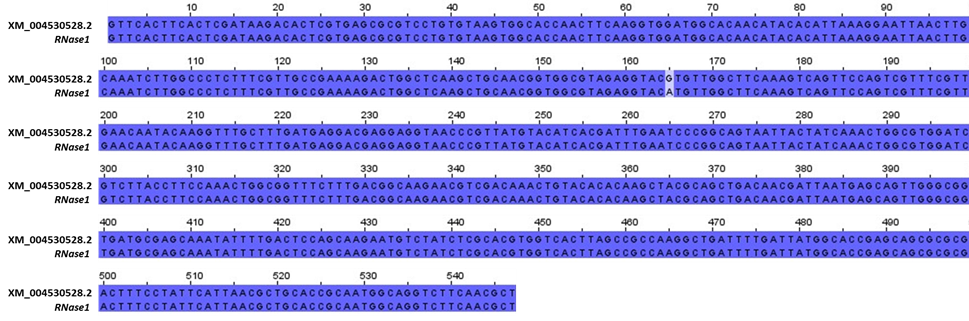

**Figure S2.** Alignment between dsRNA sequence of *RNase1 (CcdsRNase1)* gene and XM_004530528.2 (NCBI).

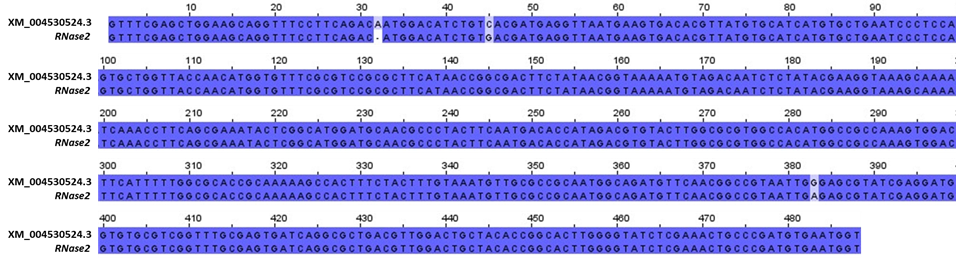

**Figure S3.** Alignment between dsRNA sequence of *RNase2 (CcdsRNase2)* gene and XM_004530524.3 (NCBI).

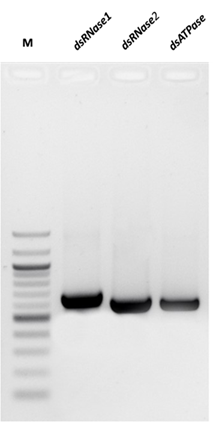

**Figure S4.** Agarose gel of the synthesized dsRNAs: dsRNase1(613 bp), dsRNase2 (557 bp), and dsATPase (553 bp).

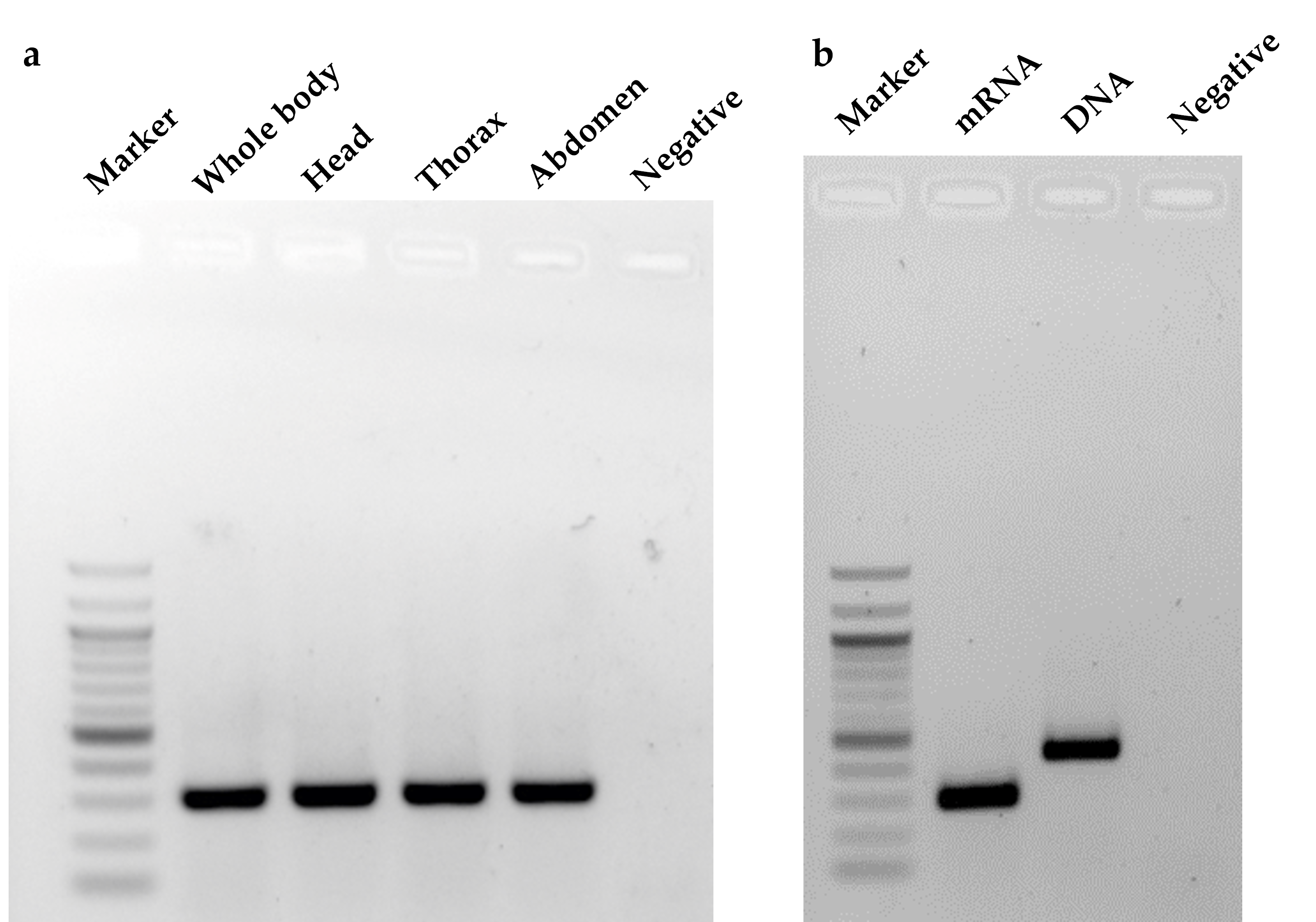

**Figure S5.** Agarose gel of *CcSOD* gene (housekeeping) expression in different adult tissues. (A) shows a 300 bp band in all samples; (B) shows the difference between the presence/absence of an intron-sequence.

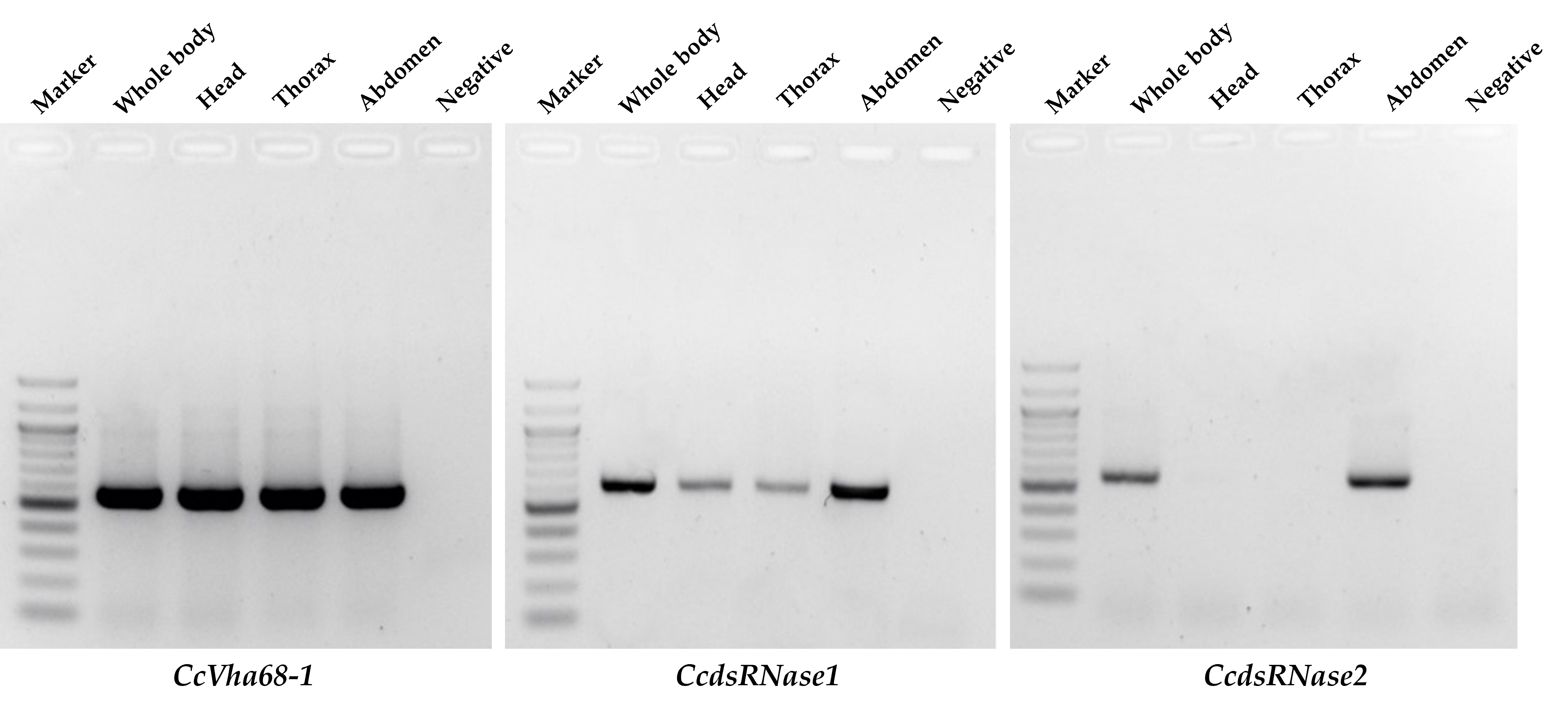

**Figure S6.** Agarose gel of *CcVha68-1*, *CcdsRNase1*, and *CcdsRNase2* gene expression in different adult tissues.

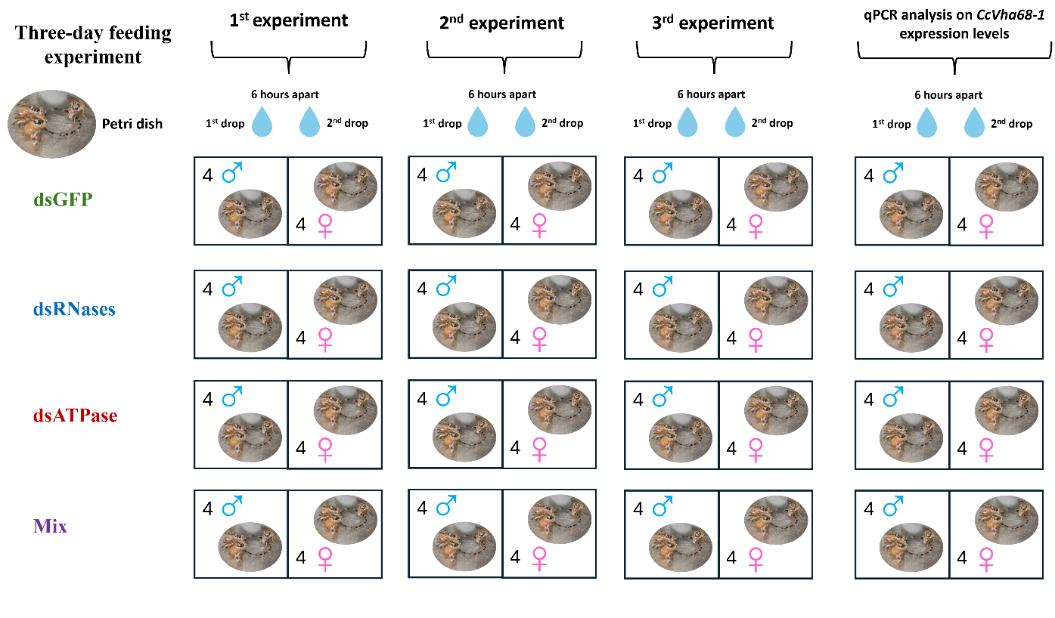

**Figure S7.** Schematic illustration of the mortality assay experiment.

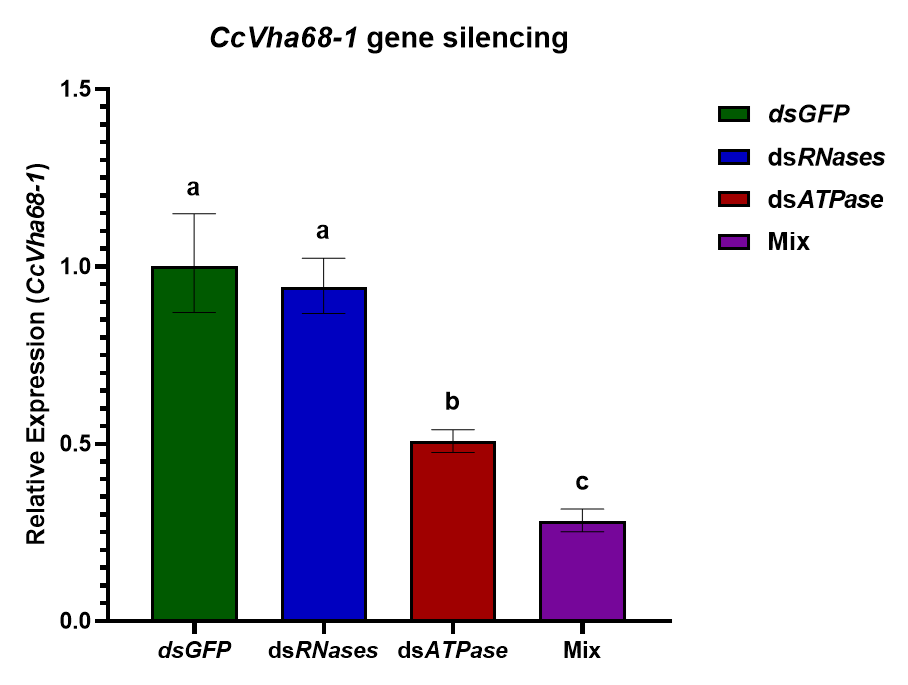

**Figure S8.** Transcript levels of *CcVha68-1* gene after three-day feeding experiment (one-way ANOVA: F(3,24) = 34.84, *P* < 0.0001). Different letters denote a significant difference between mean values recorded for each group (the obtained values passed normality tests). The values reported are the mean ± standard error.

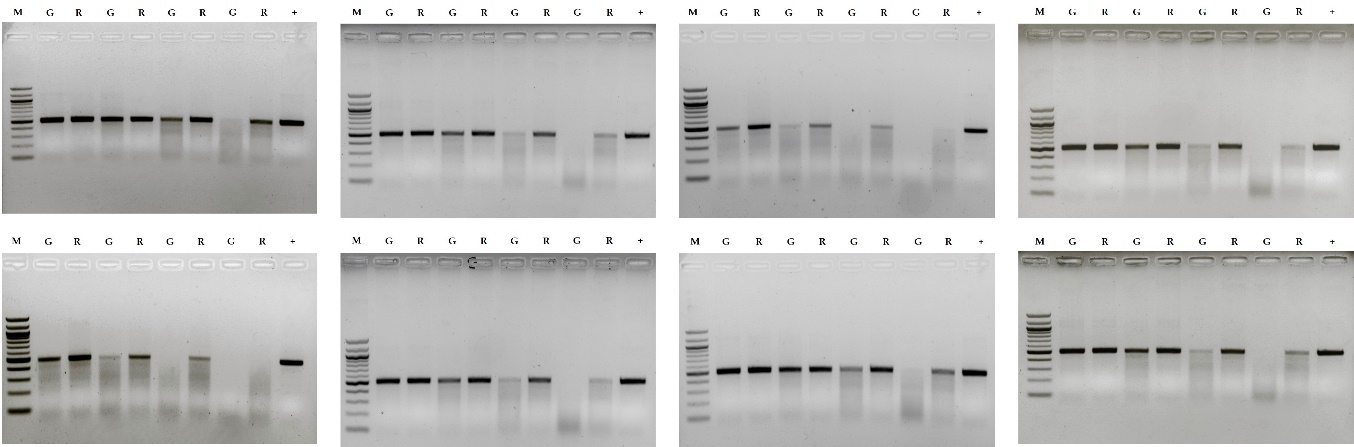

**Figure S9.** Agarose gels of *ex vivo* experiment. M: marker; G: dsGFP; R: dsRNases; +: positive control (dsATPase, 100 ng).

| Primers | Sequences | Amplicon size (bp) | Tm (°C) |
| --- | --- | --- | --- |
| **T7_dsRNase1_Fwd** | TAATACGACTCACTATAGGGAGA  GGATTTGTGGTGTCCCGGTA | 613 bp | 60 |
| **T7_dsRNase1_Rev** | TAATACGACTCACTATAGGGAGA  GCGTTGAAGACCTGCCATTG |  |  |
| **T7_dsRNase2_Fwd** | TAATACGACTCACTATAGGGAGA  CGGCACTGATCTTCTGGAGG | 557 bp | 60 |
| **T7_dsRNase2_Rev** | TAATACGACTCACTATAGGGAGA  ACCATTCACATCGGGCAGTT |  |  |
| **T7_dsATPase_Fwd** | TAATACGACTCACTATAGGGAGA  TGTTGAATTGGGTCCCGGTA | 553 bp | 60 |
| **T7_dsATPase_Rev** | TAATACGACTCACTATAGGGAGA  GTTTTGCCACAACCGAAAGC |  |  |
| **CcSOD_Fwd** | TGCTCCGAGAACGTTCACG | 300 bp | 60 |
| **CcSOD_Rev** | TCATCGGTCAATTCGTGCAC |  |  |

**Table S1.** Primers for *CcSOD* gene amplification and the synthesis of dsRNAs target.

| Primers | Sequences | Amplicon size (bp) | Tm (°C) |
| --- | --- | --- | --- |
| **RT_RNase1_Fwd** | ATTCATTAACGCTGCACCGC | 118 bp | 60 |
| **RT_RNase1_Rev** | ACCCGTCCAGCAGTCTACAT |  |  |
| **RT_RNase2_Fwd** | GGTCAAATATTCCGTGCCGC | 113 bp | 60 |
| **RT_RNase2_Rev** | TAGCTGGTGTACTCGGTCCA |  |  |
| **RT_ATPase_Fwd** | TCCCCGAACTTACTTGCGAA | 83 bp | 60 |
| **RT_ATPase_Rev** | CATGTTGGAGGTGTTGGCAA |  |  |
| **Rpl19_FWD** | AACAAACGTTGTACTGATGG | 103 bp | 60 |
| **Rpl19_REV** | CACGTACTTTATGTCGTCTG |  |  |

**Table S2.** Primers for qPCR analysis.

| dsGFP | dsRNases | dsATPase | Mix |
| --- | --- | --- | --- |
| ddH_2_O | ddH_2_O | ddH_2_O | ddH_2_O |
| Sucrose 10% | Sucrose 10% | Sucrose 10% | Sucrose 10% |
| dsGFP (200 ng/µL) | dsRNase1 (100 ng/µL) | dsATPase (200 ng/µL) | dsRNase1 (100 ng/µL) |
| / | dsRNase2 (100 ng/µL) | / | dsRNase2 (100 ng/µL) |
| / | / | / | dsATPase (200 ng/µL) |

**Table S3.** dsRNAs feeding experimental groups.
